# A Portable Fluorescence Platform for Decentralized One-Health *mcr-1* Monitoring

**DOI:** 10.64898/2026.07.24.740582

**Authors:** Maryhory Vargas-Reyes, Roberto Alcántara, Carlos Herrera, Mia Towsend, Kiara Flores-Jimenez, Carlos Raymundo, Pohl Milon

## Abstract

Antimicrobial resistance (AMR) represents a major global health threat, with plasmid-borne *mcr* genes driving colistin resistance and exposing critical gaps in One-Health surveillance across human, animal, and environmental reservoirs. The most prevalent variant, *mcr-1*, remains difficult to monitor in resource-limited settings due to the lack of rapid, affordable, and field-deployable molecular tools. Here, we developed C12a^mcr^, an integrated molecular toolbox that combines pre-amplification PCR with a fluorescent CRISPR-Cas12a assay targeting a conserved region of *mcr-1* and a custom low-cost, hand-held 3D-printed portable fluorometer. Under optimized conditions, the assay achieved a limit of detection of 630 cells/mL. In poultry feces spiked with *mcr-1*-positive *E. coli*, C12a^mcr^ detected as few as 1,800 cells/mL. When tested on 22 community-derived *E. coli* isolates, the assay showed 100% concordance with both next-generation sequencing for *mcr-1* detection and phenotypic colistin susceptibility testing by broth microdilution. The accompanying portable fluorometer performed equivalently to a laboratory microplate reader while enabling fully decentralized workflows compatible with portable PCR platforms. By integrating locally produced molecular reagents, straightforward protocols, and an accessible field-ready fluorescence reader, C12a^mcr^ overcomes key barriers to decentralized AMR surveillance and provides a practical, scalable solution for One-Health monitoring in resource-limited settings.

## 1. INTRODUCTION

Antimicrobial resistance (AMR) constitutes a global public health crisis, compromising clinical efficacy of available antimicrobial therapies. Currently, AMR is recognized as a multivariable issue involving a connection between human health and animal welfare within shared environments (Bungau et al., 2021). Resistance to colistin exemplifies the complex dynamics of AMR spread and highlights the need for a One-Health understanding to propose and implement comprehensive mitigation strategies (Bustamante et al., 2025; Rhouma et al., 2023). Such approaches should include affordable, easy-to-implement assays with competitive analytical performance for monitoring activities (Osei Sekyere, 2019). In addition, the development of portable, low-cost equipment that facilitates the transition of molecular detection assays from laboratory to field settings will support the decentralization of detection testing (Vasala et al., 2020).

Polymyxins, including colistin, were discarded for clinical use due to severe side effects, such as kidney damage and nerve toxicity (Vaara, 2019). However, the current AMR crisis reintroduced polymyxins as last-resort antibiotics for severe multidrug-resistant infections notably those caused by carbapenem-resistant Enterobacterales (El-Sayed Ahmed et al., 2020). However, their extensive prophylactic and growth-promoting use in animal farming likely precipitated widespread selection pressures and contributed to the current global dissemination of colistin resistance observed in more than 50 countries (Dadashi et al., 2022; Rhouma et al., 2023; Wang et al., 2018). Colistin resistance is primarily mediated by mobilized colistin resistance (*mcr*) variants (*mcr-1* to *mcr-10*), *mcr-1* remains the most prevalent, with 34 variants documented across accessible databases (Ling et al., 2020). Animal populations, particularly poultry and swine, serve as significant reservoirs, with reported prevalence rates ranging up to 63% to 98%, respectively (Xiaomin et al., 2020), suggesting a notable risk for zoonotic and food-chain-based horizontal resistance transmission (Nordhoff et al., 2023; Trung et al., 2017).

Currently, routine detection of colistin resistance relies primarily on phenotypic antimicrobial susceptibility testing (AST), notably broth microdilution (BMD) methods recommended by EUCAST and CLSI under ISO20776 guidelines (CLSI, 2024; EUCAST, 2025; Poirel et al., 2017). Moreover, reported molecular detection assays like LAMP, RPA, PCR-based, and whole-genome sequencing have shown high sensitivity but are still limited to laboratory setups while surveillance requires field-compatible solutions. In the current scenario, CRISPR-Cas technology has emerged as a promising alternative for molecular diagnostics, characterized by high specificity, adaptability to multiple detection formats, user-friendly protocols, and significant potential for field application (Ghouneimy et al., 2023).

Among common readouts for molecular detection assays, fluorescence-based offer superior sensitivity of benchtop microplate readers for quantitative results, restricting field applicability (Nath et al., 2023). To close this gap for fluorescence-based tests implementation, low-cost and portable fluorescence detection devices, taking advantage of 3D printing technology, have been developed and tested (Palmara et al., 2021; Rosenfeld et al., 2026; Shin et al., 2021). These portable readers aim to fulfill key design criteria, including high sensitivity, compact size, low energy consumption, and affordability (Shin et al., 2021), to effectively allow CRISPR-Cas assay implementation in decentralized and resource-limited settings.

This study addresses these challenges by proposing and evaluating a CRISPR-Cas12a-based fluorescent assay, incorporating locally produced molecular reagents and an integrated portable fluorometer. The *mcr-1* gene was selected as a model target to demonstrate the practical utility and feasibility of transitioning molecular AMR detection from centralized laboratories into potentially effective field-based detection using portable, available molecular stations.

## 2. EXPERIMENTAL SECTION

### Reagents and biological materials

Primers and crRNAs targeting a conserved region of the *mcr-1* gene were designed based on a consensus sequence for all *mcr-1* variants (*mcr-1.1* to *mcr-1.37*), retrieved from the NCBI-NDARO database (Supplementary File S1). Sequences were aligned using MEGA v.11 with default settings. For target amplification, a primer set reported by (Liu et al., 2016) was used as reference. Additionally, another set of primers was designed with Primer3Plus, excluding regions overlapping with our reference. Finally, two primer sets amplifying 309 bp (Set-01) and 268 bp (Set-02) were used in the study (Table S1).

Candidate crRNAs were predicted using CHOPCHOP (Labun et al., 2019) and CRISPRScan (Moreno-Mateos et al., 2017) tools for LbCas12a with a TTTN PAM sequence. Predicted sequences were compared and a consensus top 15 were selected for downstream analysis. Specificity was evaluated by sequence alignment against other mcr-groups, including *mcr-2*, *mcr-3*, *mcr-4*, *mcr-5*, *mcr-8*, and *mcr-9,* using MEGA v11. crRNA secondary structures were assessed using the mFOLD tool for RNA folding prediction (Zuker, 2003). Final crRNA candidates were purchased as dsDNA sequences from Macrogen Inc. (South Korea), including the T7 promoter and the scaffold sequence for LbCas12a at 5-end upstream sequence (Table S1). Designed DNA sequences’ (primers and crRNA) specificity was evaluated by sequence alignment against other mcr-group sequences (Supplementary File S2).

The crRNAs were produced by *in vitro* transcription using the TranscriptAid T7 High Yield Transcription Kit (Cat N°. K0441, ThermoFisher). The RNA products were purified with the RNA Clean & Concentrator Kit (Cat N° ZR1013, Zymo Research). Quantification was done by measuring absorbance at 260 nm in a NanoDrop One spectrophotometer. DNA Taq polymerase and LbCas12a enzymes were expressed and purified according to our previously published protocol (Mendoza-Rojas et al., 2021).

Tested *E. coli* isolates, both *mcr-1* positive and negative, were obtained from a bacterial collection generated in a previously community-based study involving children in Lima, Peru (Mendoza-Rojas et al., 2021). The study received ethical approval from the institutional review board (Approval N° 65178). Isolates were reactivated at 37°C in LB medium. Then, total DNA was purified using the Quick-DNA Miniprep plus kit (Cat N° D4069, Zymo Research). Total DNA was quantified by absorbance at 260 nm using a NanoDrop One, and stored at −20 °C.

### PCR-CRISPR/Cas12a assay (C12a^mcr^)

For target amplification and fluorescence detection, PCR and CRISPR-Cas12a reactions were performed as previously described (Vargas-Reyes et al., 2026). Briefly, 20 µL PCR reactions were prepared containing 2 ng/µL Taq DNA polymerase, 0.2 µM primers, and 1 to 10 ng/µL of DNA. Amplification was performed according to the following PCR cycling: initial denaturation at 95 °C for 3 min, followed by 25 cycles of 95 °C for 30 s, 59 °C for 30 s, and 72 °C for 30 s, and followed by a final extension at 72 °C for 5 min. PCR products were visualized in a 1.7% agarose gel stained with Trick-Track – SYBR Gold (Cat #R1161, ThermoFisher) using a Gel Doc system (Bio-Rad).

For the C12a^mcr^ reaction preparation, 150 nM crRNA was refolded (65 °C for 10 min, followed by 25°C for 10 min) and assembled with 100 nM LbCas12a, and 2 µM FAM reporter probe (6-FAM/TTATT/3IABkFQ) in CRB1 buffer (50 mM NaCl, 10 mM Tris-HCl, 100 μg/mL BSA, pH 7.9 – 25°C). The mixture was incubated at room temperature for 10 min in dark. Then, PCR products were diluted in CRB1 buffer supplemented with 20 mM MgCl₂, and mixed with the CRISPR:Cas complex to a final volume of 100 µL. Reactions were transferred to 96-well black microplates (Cat N° 23710, ThermoFisher), and fluorescence signal was recorded using a Varioskan LUX (ThermoFisher) plate multireader with an excitation wavelength at 491 nm and emission at 525.

To enhance assay performance, two variables were tested under reaction conditions identical to those previously described, except for the specific component being tested. PCR input volumes ranging from 5 to 20 µL, and alternative fluorescent probes (Table S1). Finally, the stability of the molecular components of CRISPR-Cas12a complex was evaluated over 15 days at −20 °C, 4 °C, and room temperature. Briefly, stability was estimated by comparing the fluorescent signal of positive, negative and non-template controls. A single batch of CRISPR-Cas12a complex was prepared as detailed above, and dispensed in 8 aliquots, and stored at one of the selected temperatures:-20°C, 4°C and 25°C. A freshly prepared CRISPR-Cas12a complex was used as a control. For the positive reaction, total DNA at 200 pg/µL was used. All reactions were performed using the optimized C12amcr conditions. Fluorescence was measured for 60 min using the Varioskan Lux.

### Analytical performance evaluation

The Limit of Blank (LOB) and Limit of Detection (LOD) were determined to evaluate the analytical sensitivity of the C12a^mcr^ assay. For the LOB, ten *mcr-1*-negative *E. coli* isolates were used. The LOB was calculated as: *LOB* = *Avg*_*NF_NTC_* + 1.645 *x SD* (where “Avg” is the average NF_ntc_ values, and “SD” is the standard deviation across the 10 independent reactions). The LOD was determined using both linear DNA fragments and bacterial cell counts, each tested in triplicate. For linear DNA fragments, a 1000-bp PCR product was produced using the mcr_F_1 and mcr_R_2 primers. PCR products were purified using the Oligo Clean Kit (Cat N° D4060, Zymo Research) and quantified by absorbance at 260nm. DNA concentrations ranged from 0.01 aM to 1 pM. For bacterial cell counts, an eight 4-fold dilution series, final volume of 2 mL, was prepared using a *mcr-1*-positive *E. coli* isolate. One milliliter was plated on LB agar for CFU counting, while the remaining milliliter was analyzed directly with C12a. In both cases, the LOD was then calculated using the formula *LOD* = *LOB* + 1.645 *x SD* (where “SD” corresponds to the standard deviation of the lowest analyte concentration with NF_ntc_ values above the LOB). Finally, 22 community *E. coli* isolates were evaluated using the optimized C12a^mcr^ assay. Briefly, total DNA was purified using the Quick-DNA Miniprep plus kit (Cat N° D4069, Zymo Research).

Phenotypic resistance of the evaluated *E. coli* isolates was confirmed in a broth microdilution assay. The minimum inhibitory concentration (MIC) was determined according to CLSI guidelines (CLSI, 2024). Two-fold serial dilutions (256–0.125 µg/mL) of colistin sulfate (Cat N° C4461-100 MG, Sigma Aldrich) were prepared in Mueller– Hinton Broth II (Cat N° 610218, Liofilchem). Cell suspensions at 1.5 × 10^6^ CFU/mL were inoculated, including *E. coli* ATCC 25922 as a control. After 18 h of incubation at 37 °C, the MIC was determined as the lowest colistin concentration that inhibited visible bacterial growth.

### Portable fluorometer design

The portable fluorometer was designed and manufactured to enable the detection of fluorescence signals compatible with portable molecular biology stations, such as the Bentolab. The fluorometer is currently under patent application with INDECOPI, Peru, with ID number 003127-2024/DIN. (Herrera *et al.,* 2024). The fluorometer was modeled using SolidWorks computer-aided design software and subsequently fabricated using a Prusa MK4S 3D printer, based on the filament deposition process. The printed components of the device structure were fabricated using black and white polylactic acid (PLA) (eSUN, China): black was used for the internal optical path to minimize light interference, while white was used for the external casing. The printed circuit boards (PCBs) were machined with an xsTECH CNC milling machine, and the electronic components were subsequently assembled. Fluorescence values are obtained by evaluation of a region of interest (ROI1) defined in a digital image of the measured reaction tube. From ROI1, the histogram of the green channel is obtained, and the weighted average intensity is calculated using the values of the histogram. This value is considered the fluorescence intensity associated with the analyzed region.

To determine the positivity of a sample, a non-template control (NTC) is first measured to obtain a reference intensity value. This value defines a selectable threshold, equivalent to 1.5-or 2-fold of the NTC fluorescent intensity. Subsequently, the test samples are measured and, if the calculated intensity exceeds the threshold, the sample is classified as positive; otherwise, it is classified as negative. Additionally, a graphical user interface was developed in Python using the Tkinter and Openpyxl libraries.

### Portable fluorometer performance evaluation

Signal stability was evaluated directly using C12a^mcr^ using positive, negative, and NTC controls in the portable fluorometer at 10-minute intervals over 100 min to define an appropriate reading window. Subsequently, target sensitivity was evaluated using a DNA concentration curve ranging from 10³ to 10⁸ ag/µL. Fluorescence measurement was compared between the portable fluorometer and the Varioskan Lux fluorescence reader (ThermoFisher) as reference. Fluorescence signals were monitored each 10 minutes for 1 hour. Then, three representative DNA levels were selected for comparing the signal threshold for target detection: high (200 pg/µL), medium (20 pg/µL), and low (0.2 pg/µL). Fluorescence was measured in the portable fluorometer at 20, 30 and 40 minutes of the CRISPR reaction. All indicated assays were done in triplicate.

Finally, fecal mock samples were prepared by spiking fecal suspensions as previously reported (Naas et al., 2011) to evaluate the target detection in simulated conditions. Briefly, fecal samples were obtained from a poultry farm dedicated to chicken production for human consumption in Quilmaná, Lurín, Peru. A fecal suspension was prepared by dissolving 6 g of feces in 60 mL of sterile water. An *E. coli mcr-1* positive isolate was then spiked at three bacterial concentrations ranging from 1.8 x 10^3^ to 1.8 x 10^5^ CFU/mL. The negative control consisted of fecal samples without bacterial cells.

Total DNA was extracted using the ZymoBIOMICS DNA Miniprep Kit (Cat N° D4300, Zymo Research) according to the manufacturer’s instructions. Then, PCR amplification was performed as detailed above in the portable workstation BentoLab, in a 20 μL reaction. After amplification, the C12a^mcr^ reaction was prepared as described above and added directly to the PCR tube to a final volume of 100 μL. After 20 min incubation in the dark, the fluorescence signal was detected using the portable fluorometer device.

## 3. RESULTS

### C12a^mcr^ optimization

Before evaluating the fluorometer performance, a CRISPR-Cas-based assay was developed and optimized for detecting *mcr-1* genes (C12a^mcr^ assay) in isolated *E. coli* strains. Two primers sets and three crRNA sequences were selected for targeting two different regions in the *mcr-1* gene (Fig. S1A). Both primer sets showed specific target amplification as confirmed by visualizing unique PCR bands for positive controls.

Detection primer Set-01 produced a 309-bp PCR product, while Set-02 produced a 268-bp amplicon. No differences in target amplification efficiency were observed (Fig. S1B). Nevertheless, coupling the primers Set-01 with the crRNA-1b resulted in the highest fluorescence signals in comparison to crRNA-1a and the full Set-02, with more than 5-fold higher than the no-template control (NTC) up to 2.5 pg/µL of template (Fig. 1A).

**Figure 1.**
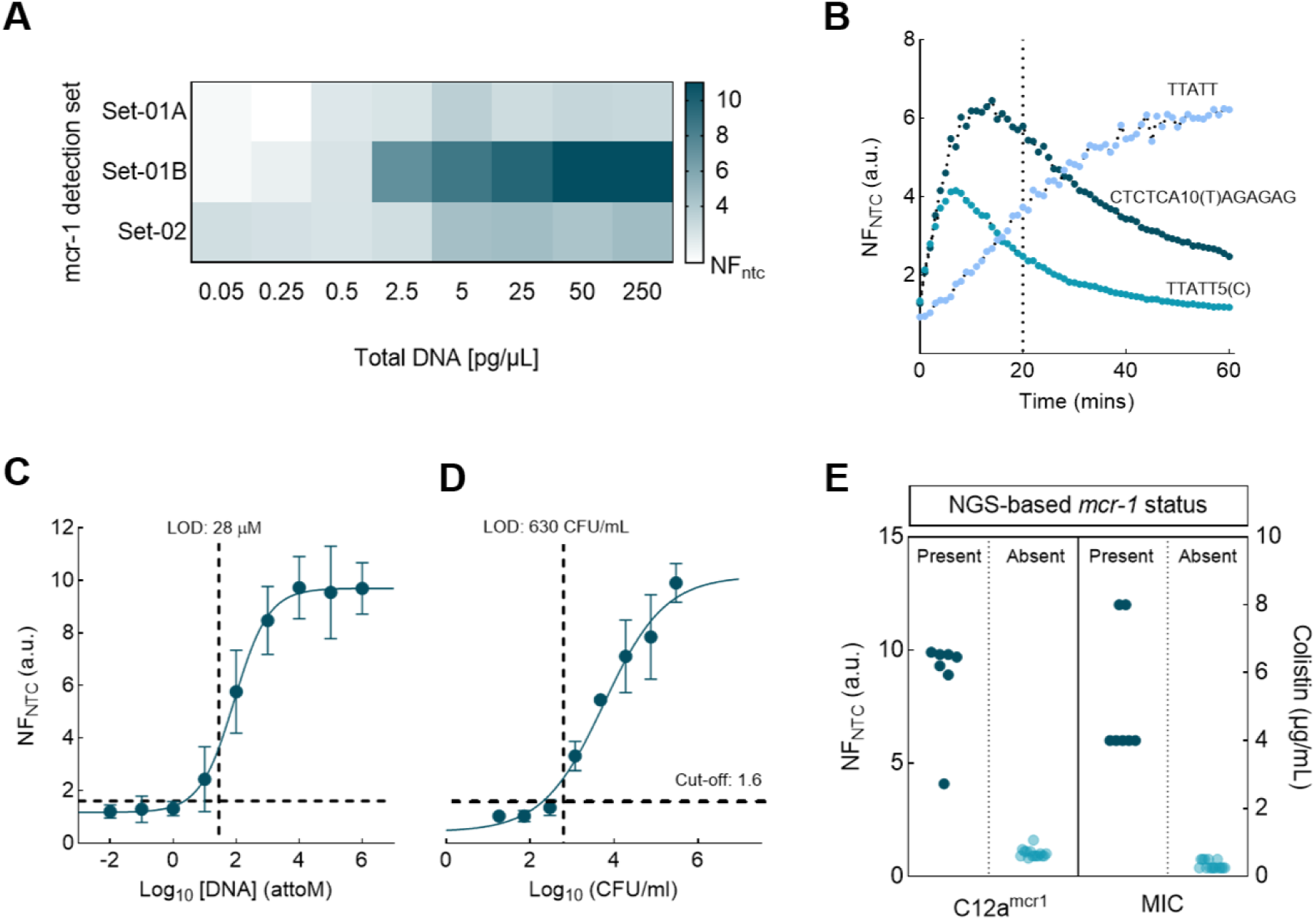
Fluorescence-based analytical performance of a C12a^mcr^ for the detection of colistin-resistance *mcr-1* gene. **(A)** Detection performance of primers and crRNA sets (Table S1) was evaluated across total DNA concentrations ranging from 0.05 to 250 pg/µL. Normalized fluorescence relative to the non-template control (NF_ntc_) calculated at 20-min is shown. **(B)** Fluorescence signal as a function of different labeled probes. NF_ntc_ values were calculated over 60-min. The optimized time for the assay is indicated by the dotted line at 20-min. Probes sequences: CTCTCA10(T)AGAGAG (Hairpin), TTATT and the TTATT5(C). Limit of detection (LOD) estimated for **(C)** using linear DNA target sequences and for **(D)** cell number (CFU) per mL. Dotted line on the Y-axis represents the Limit of Blank (LOB) estimated for the C12a^mcr^ assay (LOB = 1.6). Dotted line on the X-axis marks the calculated LOD at a 20-min reaction. **(E)** C12a^mcr^ detection using 22 isolated *E. coli* strains, previously characterized by NGS with *mcr-1* present (n = 7) or absent (n = 15). The left-side graph shows the NF_ntc_ values from the C12a^mcr^ assay. The right-side graph presents the Minimal Inhibition Concentration (MIC) assay using 2 µg/mL colistin as threshold (CLSI, 2024). Dark cyan points represent resistant/*mcr-1* positive strains, while light cyan points represent susceptible/negative *mcr-1* strains.

Based on this screening, the combination of the primer Set-01 with the crRNA-1b was selected for downstream optimization. To increase fluorescence signal for low target concentrations, we evaluated four DNA template concentrations (ranging 2.5 x 10^-2^ to 5 x 10^-1^ pg/µL) with a 25 and 30-cycles PCR. An increase of five amplification cycles produced a 4-fold higher signal for the lowest tested DNA target concentration. Similarly, a 6-fold increase in fluorescence was observed for the highest tested concentration, in comparison to the 25-cycle PCR input (Fig. S2A). One technical limitation of many CRISPR-Cas-based assays is that amplification and detection must be performed separately because the required enzymes have different thermal tolerances, especially the DNA Taq polymerase and LbCas12a. Therefore, an aliquot of the amplification reaction is typically transferred into the CRISPR-Cas detection reaction for signal generation. Here, we evaluate whether reversing the order of addition (i.e., introducing the CRISPR-Cas reaction into the full PCR volume rather than transferring an aliquot from PCR) can enhance assay practicality and facilitate kit-like formats.

Different PCR reaction volumes were tested as CRISPR-Cas reaction input, including the whole PCR volume (20 µL). Regardless of DNA template concentration, higher PCR volume produced higher fluorescent signals. This effect was better observed for the lower tested target concentrations, reaching 2.5-fold higher NF_ntc_ between 5 and 20 µL PCR input. No differences were observed for the negative control (1.47 ± 0.18 a.u.), suggesting the background remained stable (Fig.S2B).

Additionally, different probes, varying in length and secondary structure, were evaluated at 200 nM final concentration. The hairpin-containing probe showed the highest NF_ntc_ value at a 20-min cutoff (Fig. 1B). However, the canonical probe (TTATT) showed a more stable background as the normalized fluorescence signal showed a steady increase over time compared to the other tested probes (Fig. 1B).

Finally, to support the intended field application of the assay and instrument, the stability of the molecular components was evaluated. Although lyophilization is the standard method for preserving enzymes without a cold chain, its use can be limited by access to specialized equipment. Therefore, we tested if the crRNA:Cas12a complex can tolerate storage at common temperatures commonly available in basic laboratory settings (4°C and-20°C) plus room temperature (∼25°C). The stability was tested using a DNA target concentration of 200 pg/µL of. C12a^mcr^ produced fluorescence across 15-days of incubation period. However, signal intensity declined over time, reaching an average 50% loss across all storage temperatures by day 15. Storage at-20°C preserved the crRNA: Cas12a complex better, with less than 30% signal loss up to day 3 (Fig. S3A).

In contrast, assay specificity remained up to 8 days of incubation under all conditions. At-20°C, no false-positive results were observed during the entire 15-day evaluation period (Fig. S3B).

### C12a^mcr^ performance

Based on the optimized assays, the final conditions for the C12a^mcr^ assay included 20 µL PCR input, using the hairpin-containing probe with a 20-min reading cutoff. Analytical sensitivity was evaluated through LOB and LOD determination. At 20-min incubation, a LOB of 1.6 was estimated. Background signal was kept relatively constant over a 60-min reading (NF_ntc_ mean of 1.15 ± 0.07 a.u.) (Fig. S2C). Under optimized reaction conditions, a LOD of 28 aM was estimated using a 1000-bp linear DNA sequence as target; meanwhile, a LOD of 630 cells/mL was determined for cell counts. In both cases, the lowest detectable target concentration showed a NF_ntc_ value of more than two (Fig. 1C-D). The optimized assay was used with 22 isolated *E. coli* strains that were previously characterized by NGS sequencing to confirm *mcr-1* gene presence. The 22 isolates were evaluated by AST to determine their colistin susceptibility profile. Perfect agreement between phenotypic profiling, NGS-*mcr-1* gene presence and C12a^mcr^ detection was observed (Fig. 1E, Table S2). Positive-*mcr-1* isolates showed NF_ntc_ values more than 5 and MIC more than 5 µg/mL.

### Portable fluorometer hardware and software functionality

The portable fluorescence reader developed in this study is a compact, hand-sized device designed for quantitative detection of the fluorescent signal generated by the C12amcr assay (Fig. 2A, Fig. S4A-B). The instrument was fabricated de novo using 3D-printed components (Prusa MK4S printer, black and white PLA filament) at an estimated total cost of approximately 140 USD, including all electronic and optical parts (Table S3). It features a light-tight detection chamber with a magnetic lid that accommodates standard 0.2 mL PCR tubes containing 0.1 mL of the reaction, enabling direct transfer of tubes from a thermocycler (Fig. 2A). Excitation and detection follow a 90° right-angle geometry: a high-power XLamp XQ-E Plus LED (peak emission 475 nm) illuminates the sample from below, while emitted fluorescence passes through a 530 nm long-pass filter and is captured by an RGB color sensor. The optical path is housed in a low-reflectance tunnel to minimize stray light (Fig. 2B).

**Figure 2.**
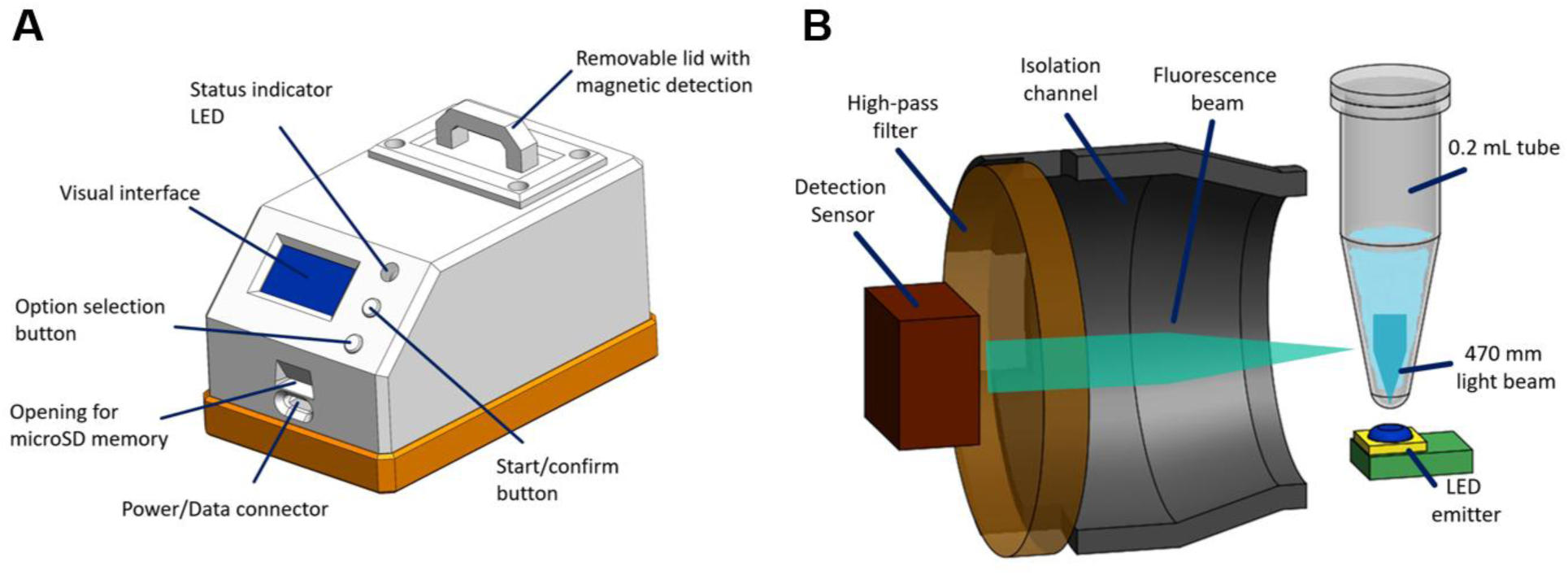
Hardware fluorometer design. **(A)** Schematic of the fluorometer displaying the main components. The device uses a microSD card to store the generated signal data. Power is supplied by a portable power bank (5V, ≥ 1 Amp, with low current mode option) or through direct connection to a regular laptop. The visual interface displays the detection result as “POS” (for positive) and “NEG” (for negative). **(B)** Core components scheme displaying the detection compartment for fluorescence measurement of a 0.2 mL polypropylene tube (i.e., as commonly used for PCR) containing 100 µL reaction. The LED emitter is positioned beneath the tube holder, while emitted fluorescence is filtered through a high-pass filter placed perpendicularly to the excitation beam before reaching the detection sensor.

Image acquisition and analysis are performed automatically. The green channel of the captured image is extracted, and a fixed region of interest (ROI1) corresponding to the full 100 µL reaction volume is defined. Fluorescence intensity is quantified as the weighted average of the green-channel histogram values and reported in arbitrary units (a.u.). Sample classification uses a negative-control baseline; a configurable threshold of 1.5-or 2-fold of this baseline intensity determines a positive or negative result. The device includes an integrated OLED screen that displays real-time fluorescence values, user settings, and the detection outcome (POS/NEG/INVALID), as well as a microSD card slot for automatic storage of raw tube images (Fig. 2A). A dedicated graphical user interface developed in Python (Tkinter and Openpyxl libraries, packaged as a standalone.exe with PyInstaller) enables visualization of images, reference levels, calculated thresholds, and one-click export of all data as XLS files (Fig. S4C). The reader operates on standard 5 V power banks (≥ 1 A) or laptop USB

### Portable fluorometer performance characterization

Once the C12a^mcr^ assay was optimized, we evaluated the designed fluorometer for detecting fluorescence signals using a portable operational setup including portable workstations like BentoLab (Fig. S4B). For fluorescence detection, the CRISPR-Cas reaction mix was added to the 20 µL PCR, incubated for 20 min at room temperature, and then measured in the portable fluorometer. Positive reactions showed fluorescence across the whole reaction volume compared to negative reactions (Fig. 3A). Negative and non-template control reactions showed fluorescence lower than 50 a.u., while positive reactions showed 2-fold higher fluorescence on average (Fig. 3B).

**Figure 3.**
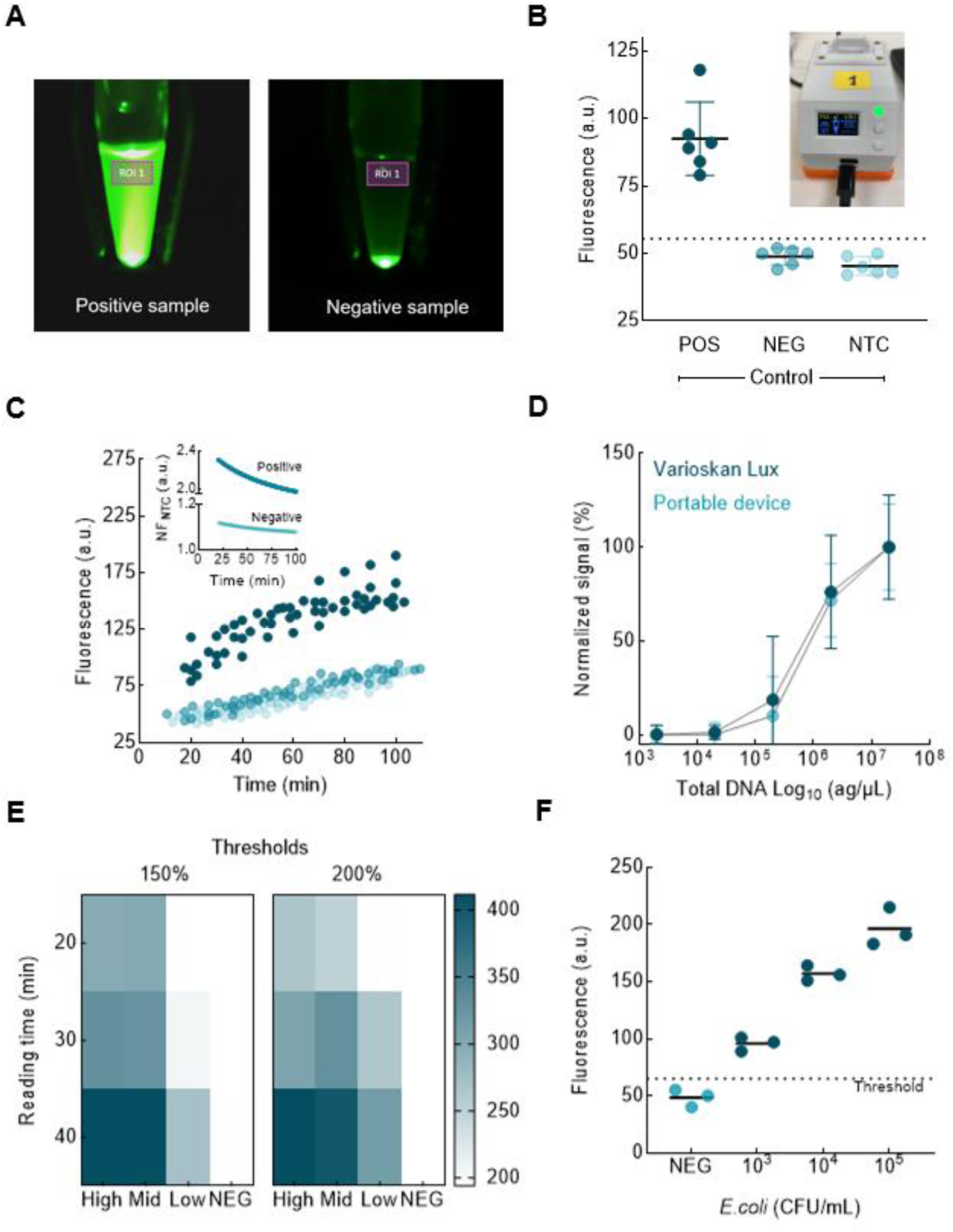
C12a^mcr^ performance with the portable fluorescence reader. Fluorescence measurements in the portable fluorometer were performed using the optimized C12a^mcr^ assay. **(A)** Example images captured by the portable fluorometer for positive and negative samples. Image analysis is performed within a fixed region of interest (ROI), independent of the sample type. **(B)** Fluorescence measurements from a set of samples (n = 6) including positive-, negative and non-template controls (NTC). The dotted line in the Y-axis represents the mean plus three standard deviations of the average NTC fluorescence signal. The inset shows the fluorometer working while measuring fluorescence. **(C)** Fluorescence measurements collected over a 100 min for positive (dark cyan), negative (cyan) and non-template (light cyan) reactions. Each time-point represents six replicates displayed with staggered scattering. The inset plot shows the normalized fluorescence ratio (NF_ntc_) for positive and negative reactions relative to the non-template control (NTC). **(D)** Comparison of fluorescence measurements obtained using the portable fluorometer and the Varioskan Lux microplate reader (ThermoFisher) across DNA concentrations ranging from 2 fg/µL to 20 pg/µL. Fluorescence values were normalized to a percentage for better comparison. Measurements were performed in triplicate. **(E)** Performance comparison of detection thresholds in detecting three target concentrations Low (0.2 pg/µL), Mid (20 pg/µL), and High (200 pg/µL). Fluorescence values (a.u.) are shown for 20, 30 and 40 min. **(F)** C12a^mcr^ detection performance using mock poultry feces samples spiked with three serial 10-fold dilutions of a *mcr-1*-positive *E. coli* strain. Measurements were performed in triplicate. Raw fluorescence values (a.u.) are plotted, and the black line in each group represents the mean fluorescence. The dotted line in the Y-axis represents the threshold value based on the non-template control. The evaluated bacteria dilutions correspond to final concentrations of approximately 1.8 × 10³, 1.8 × 10⁴, and 1.8 × 10⁵ cells/mL.

Fluorescence signal was measured for 60 min. Negative and non-template control did not show large differences over time; by contrast, positive reactions showed 2-fold higher fluorescence compared to negative or NTCs during the entire time course (Fig. 3C). Normalized fluorescence showed that positive and negative reactions were discriminated along the recording time, although unspecific signals also increased over time (Fig. 3C-inset).

Then the portable fluorometer was compared with a benchtop instrument, the Varioskan Lux (ThermoFisher), by measuring fluorescence in the optimized conditions using a DNA target dilution curve ranging from 2 fg/µL to 20 pg/µL. Fluorescence was normalized to a percentage for better comparison. The results indicate that the portable reader can detect similar DNA concentrations as the commercial plate reader, as no differences were observed in the analytical sensitivity of the C12a^mcr^ assay (Fig. 3D).

Two fluorescence thresholds were compared for the portable fluorometer. These cutoffs represent the minimum fluorescence signal (a.u.) required for a sample to be classified as positive and are defined as 1.5-and 2-fold of the fluorescence value measured for the NTC reaction. Three C12a^mcr^ reactions with different DNA template concentrations were measured, 0.2 pg/µL (low), 20 pg/µL (medium) and 200 pg/µL (high) at three reading times. At the 1.5-fold cutoff, all target concentrations were detected confidently at 40-min incubation. In contrast to the 2-fold cutoff, for which all concentrations crossed the threshold at 30-min. Only the low target concentration was not detected at the 20-min reading independently of the selected threshold (Fig. 2E). Finally, mock poultry feces samples were assessed. A positive trend was observed between the fluorescence signal and cell count. As observed, 1800 cells/mL were detected using C12a^mcr^ with the portable fluorometer (Fig. 2F). The sample matrix did not show fluorescence higher than the 2-fold threshold.

## 4. DISCUSSION

Addressing the global spread of antibiotic resistance gene (ARG) requires comprehensive surveillance across multiple interconnected environments. Affordable and competitive detection tools, including both diagnostic assays and instrumentation, are therefore essential, especially in low-resource settings. Access to 3D-printing and open-source strategies has boosted the development of low-cost, user-friendly, and technically compliant instruments that perform on par with benchtop devices (McNair et al., 2024). In this study, we optimized a CRISPR-Cas-based assay to detect the colistin-resistance-associated *mcr-1* gene with a 3D-printed, hand-size fluorometer.

Among *mcr* family members, *mcr-1* gene is the most prevalent (Ling et al., 2020), with worldwide dissemination, possibly aided by food-chain transmission. Multiple animal reservoirs, including poultry and livestock, have been reported to carry *mcr-1*-harbouring *Enterobacteriaceae* (Wang et al., 2018; Xiaomin et al., 2020). Conventional antimicrobial susceptibility testing is time-consuming for continuous monitoring (Vasala et al., 2020), and while molecular assays for detecting *mcr* genes have been reported (Osei Sekyere, 2019), most rely on standard laboratory infrastructure. CRISPR-Cas-based assays offer a novel, highly sensitive alternative with potential for POC-like approaches (Ghouneimy et al., 2023). Our C12a^mcr^ assay detected as low as 1.8 x 10^3^ CFU/mL in spiked fecal samples within a 20-min window detection. For *mcr-1* detection, previous nucleic-acid amplification-based assays reported sensitivities ranging 6.3×10^3^ to 10^4^ CFU/mL for PCR (Gong et al., 2021, 2019; Zou et al., 2017). Compared with traditional molecular approaches, our C12a^mcr^ assay offers competitive sensitivity without requiring qPCR instruments or electrophoresis for readout visualization.

CRISPR-Cas assays have been reported for several ARG (Demirayak and Akay Sazaklioglu, 2026; Quiroz-Huanca et al., 2026; Vargas-Reyes et al., 2026), demonstrating competitive performance with sensitivities ranging 7.5 x 10^-1^ fg/µL to 5 x 10^3^ fg/µL for genomic DNA from either clinical and environmental samples (Li et al., 2024; Mao et al., 2024). Regarding cell counts, a sensitivity of 10^5^ cells/mL has been reported (Gao et al., 2025). For *mcr-1*, specifically, two studies have described the use of RPA combined with CRISPR-Cas detection. One study achieved a sensitivity of 6.2 x 10^3^ cells/mL in spiked clinical stool samples, with a 60-min turnaround time (Gong et al., 2022). Similarly, using the AacCas12b ortholog yields a sensitivity of 7 CFU/g in spiked pork meat samples, with a total reaction time of 55 min (Wang et al., 2024). The analytical sensitivity of our C12a^mcr^ assay is comparable when considering similar sample types (i.e., stool). Although those studies reported shorter turnaround times (< 60 min) than our proposed workflow (∼100 min) (Tan et al., 2022). In contrast, Taq DNA polymerase continues to be the most widely used enzyme for nucleic acid amplification assays and, as reported here, unlocks the implementation of molecular assays using entirely locally produced molecular reagents (Mendoza-Rojas et al., 2021).

Various readout formats have been reported for CRISPR-Cas assays, including colorimetric, fluorescent and electrochemical detection (El-Sayed Ahmed et al., 2020). Fluorescence offers a more favorable signal-to-noise ratio, providing 10 – 100-fold higher sensitivity (Zheng et al., 2023) but requires a dedicated reader, making it a major limitation for implementation outside laboratory settings (Lau et al., 2025; Nath et al., 2023). Portable readers should therefore offer high sensitivity, small size, low power consumption, and affordability to be competitive (Katzmeier et al., 2019; McNair et al., 2024; Shin et al., 2021). In this study, the designed fluorescence reader demonstrated sensitivity comparable to benchtop equipment, such as the Varioskan Lux. Similarly, several locally designed fluorescence readers have also shown performance levels like standard laboratory instruments (Katzmeier et al., 2019; McNair et al., 2024; Rosenfeld et al., 2026), supporting the feasibility of integrating low-cost devices into POC workflows.

Although benchtop readers provide automation and analytical versatility, their complexity and cost limit use in resource-limited settings. In contrast, portable instruments stand out for their low operational complexity, and reduced cost, offering an accessible alternative that retains key quantitative advantages without requiring technical complexity. Our portable reader was designed for hand-size portability to facilitate transport similar to Nanopore devices (Dai et al., 2025; Katzmeier et al., 2019; Quero et al., 2025), and is compatible with common power banks or direct laptop connection, making it ideal for field applications (Dai et al., 2025; Quero et al., 2025; Rosenfeld et al., 2026). While throughput is lower than that of plate-based instruments, the portability-to-performance trade-off is favorable for surveillance and screening use cases.

A technical comparison between the portable fluorometer, a benchtop multireader, and a blue-light transilluminator (Table S4) showed that the portable reader provides an intermediate solution as it retains portability and low cost like qualitative visual devices while offering integrated digital fluorescence processing closer to quantitative benchtop instruments. The designed portable instrument combines image-based fluorescence detection with integrated digital processing, enabling quantitative measurement that is one of the principal limitations of visualization-dedicated instruments.

A persistent challenge for portability integration is carryover contamination (Lau et al., 2025), especially when amplification and detection occur in separate steps (Zu et al., 2025). One-tube CRISPR-Cas formats coupled to isothermal amplification have used physical separation (Shi et al., 2022), chemical additives (Li et al., 2021) to mitigate sensitivity loss. However, these approaches are not compatible with PCR-based amplification. Here, we implement the concept of using CRISPR-Cas reaction analogously to standard detection-kit reagents, by adding CRISPR-Cas mixture directly to the amplification reaction. This approach maintained strong fluorescence signals even at high target concentrations while reducing contamination risk and simplifying the protocol.

Reagent stability is another challenge for the implementation of molecular assays outside laboratory settings (Ghouneimy et al., 2023). For CRISPR-Cas-based assays, preserving Cas proteins and crRNA integrity is critical. Although lyophilization, filter-drying and chemical stabilization have been explored (Li et al., 2021; Rybnicky et al., 2022; Sen et al., 2024), they are not easy to implement locally. Cold-chain storage remains the most practical method for preserving reagent stability, and basic laboratory refrigeration is often accessible even in minimally equipped settings. We observed that the Cas:crRNA complex retains activity with basic cold storage, particularly at –20 °C, which is achievable with standard freezers. Further assessment should optimize stabilization strategies for prolonged field-based monitoring activities.

## 5. CONCLUSION

In conclusion, we have developed and validated C12a^mcr^, a complete CRISPR-Cas12a-based fluorescent assay integrated with a low-cost, 3D-printed portable fluorometer for decentralized detection of the colistin-resistance gene *mcr-1*. The optimized workflow achieved strong analytical performance, with limits of detection of 630 cells/mL in pure culture and 1,800 cells/mL in complex poultry feces matrices, and delivered 100% concordance with gold-standard next-generation sequencing and phenotypic antimicrobial susceptibility testing across 22 E. coli isolates. The hand-held fluorometer (∼US$140 total cost) matched the performance of benchtop instruments while remaining fully compatible with portable thermocyclers, enabling molecular AMR surveillance outside centralized laboratories. By relying on locally produced reagents and simple protocols, this platform directly addresses accessibility barriers in resource-limited settings and advances practical One-Health strategies against emerging resistance threats. Future efforts will focus on real-world field validation, potential multiplexing for additional *mcr* variants or antimicrobial resistance genes, and improved reagent stabilization to minimize cold-chain requirements. C12a^mcr^ offers a scalable model for democratizing molecular diagnostics in the global response to antimicrobial resistance.

## DECLARATIONS

## Acknowledgement

We are very thankful to Dr. Mónica Pajuelo for providing access to the *E. coli* isolate sample set used in this study.

## Ethics

The *E. coli* isolates used in this study were anonymous and derived from a biorepository of a longitudinal study approved by the ethics committee of Universidad Peruana de Cayetano Heredia (UPCH, SIDISI: 65178).

## Declaration of generative AI in scientific writing

During the preparation of this work, the authors used AI tools to improve readability and for checking grammar and spelling. After using this tool, the authors reviewed and edited the content as needed and took full responsibility for the content of the publication

## Sequence information

No datasets were generated during the current study.

## Author contribution

**R. Alcántara:** Conceptualization, methodology, formal analysis, visualization, writing-original draft, supervision, funding acquisition, writing-review & editing

**M. Vargas-Reyes**: Investigation, formal analysis, writing-original draft

**K. Flores:** Investigation

**C. Herrera:** Investigation, formal analysis, software, visualization, writing-original draft

**M. Townsend:** Investigation, formal analysis, visualization

**C. Raymundo:** Supervision, methodology, funding acquisition, writing-review & editing

**P. Milón:** Conceptualization, supervision, funding acquisition, writing-original draft, writing-review & editing

